# An epigenetic clock for *Xenopus tropicalis* reveals age-associated sites enriched in H4K20me3-marked heterochromatin

**DOI:** 10.1101/2024.12.21.629929

**Authors:** Ronan Bennett, Marco Morselli, Leonid Peshkin, Matteo Pellegrini

## Abstract

DNA methylation clocks have been widely used for accurate age prediction, but most studies have been carried out on mammals. Here we present a clock for the aquatic frog *Xenopus tropicalis*, a widely used model organism in developmental biology and genomics. The clock, trained on targeted bisulfite sequencing data, achieved a cross-validation median absolute error of 0.70 years. To construct the clock, we collected DNA methylation data from 192 frogs at genomic regions containing CpG sites previously shown to have age-associated methylation in *Xenopus*. We found that CpG sites with methylation levels significantly increasing or decreasing with age are enriched in heterochromatic regions marked with H4K20me3 and H3K9me3. CpG sites with positively age-associated methylation levels are enriched in bivalent chromatin and gene bodies with H3K36me3, and tend to be proximal to lowly expressed genes. These epigenetic features of aging are similar to those found in mammals, suggesting evolutionary conservation of epigenetic aging mechanisms. Our clock enables future aging biology experiments that leverage the unique properties of amphibians.

## Introduction

DNA methylation clocks have been broadly used for age prediction in mammals [1]. A common approach for constructing these clocks is to use a regularized linear regression model to predict chronological age based on DNA methylation levels. Certain CpG sites have methylation levels that reliably increase or decrease with age, so these linear clocks are able to accurately predict age based on these sites. These clocks allow the determination of the age of animals caught in the wild, which opens up new research opportunities in conservation biology.

Another common application of epigenetic clocks is to determine the rate of aging of an individual, by measuring whether they are aging faster or slower than other individuals with the same chronological age. This property is useful for studies that evaluate the impact of interventions on lifespan. For long-lived species, an experiment that directly measures the time until death can be prohibitively long, but an epigenetic clock can provide a more immediate endpoint to evaluate the effectiveness of an intervention, thus accelerating the pace of aging research. In humans, epigenetic age acceleration (the number of years that epigenetic predicted age exceeds chronological age) is correlated with all-cause mortality and a number of major diseases such as cancer and heart disease [2–4]. The predicted age from DNA methylation clocks, also known as epigenetic age, has been shown to slow down with interventions that extend lifespan in mice such as caloric restriction [5]. Epigenetic clocks evaluated in human fibroblasts before and after treatment with Yamanaka factors to obtain induced pluripotent stem cells showed epigenetic age decrease to near zero after the treatment, similar to the epigenetic age of other stem cells [1,5]. Furthermore, partial reprogramming of somatic cells induced a steady decline in epigenetic age over 20 days of OSKM expression until it reached zero [6].

People with diseases of accelerated aging such as Werner syndrome and Hutchinson-Gilford progeria syndrome have epigenetic ages that are significantly higher than controls [7,8]. These observations suggest that DNA methylation clocks can measure cellular aspects of the biological aging process. However, the mechanisms underlying age-associated methylation changes are poorly understood. Age-correlated CpG sites have been preferentially found in loci that are targets of PRC2 [9,10], and near promoters of transcription factors involved in developmental processes in humans [9,11] and *Xenopus* [12].

While the majority of the previous studies of epigenetic aging have been carried out in mammals, clocks have also been developed for other vertebrates, including amphibians. Amphibians are invaluable models for biological research due to their unique life cycles, diverse ecological niches, and physiological adaptability. Their ability to live in both aquatic and terrestrial environments provides insights into developmental biology, environmental adaptation, and evolutionary processes. Amphibians, like frogs and salamanders, are particularly useful in studying regeneration, as many can regenerate limbs and organs, offering models for tissue repair and stem cell research. Additionally, their permeable skin and sensitivity to environmental changes make them excellent indicators for ecological and toxicological studies, helping researchers understand the impact of environmental stressors on living organisms. An amphibian epigenetic clock could advance aging research by providing a tool to measure biological age with precision in species that exhibit unique regenerative abilities and variable lifespans. By studying how epigenetic markers correlate with aging in amphibians, researchers may uncover mechanisms of longevity, regeneration, and cellular resilience, offering insights that could inform therapies for aging and age-related diseases in humans.

An example of an amphibian epigenetic clock is the study by Zoller et al. [12] which constructed DNA methylation clocks in *Xenopus tropicalis* and *Xenopus laevis*. This study used the mammalian methylation array [13] to measure the methylation state of 4,635 CpG sites that have high sequence conservation between *Xenopus* and mammals. They demonstrated evolutionary conservation in DNA methylation changes between *Xenopus* frogs and humans by constructing joint human-frog clocks that are able to predict the age of either species. This supports the use of *Xenopus* as a model organism to study epigenetic aging. While this dataset is well-suited for studying similarities in DNA methylation levels between *Xenopus* and mammals, it is limited by the fact that the use of the mammalian methylation array excludes the vast majority of CpG sites where methylation changes are specific to *Xenopus*. Moreover, the study is also limited in the number and type of samples used. They reported samples from 6 tissues pooled from a number of individual frogs (35 *X. laevis* and 30 *X. tropicalis* tissue samples).

However, they encountered technical challenges that resulted in the loss of over half of the samples [14]. Thus, while this study represents a promising first step, because of the above limitations it is unlikely that the resulting clock or the mammalian array will be broadly used for aging studies in frogs.

In the present work, we sought to construct a robust DNA methylation clock based on data from 192 frogs, focusing on a single easily sampled tissue, and measuring a comprehensive set of *Xenopus*-specific age-associated CpGs. We generated methylation data from *Xenopus tropicalis* frogs using a skin punch from the hindlimb webbing, followed by DNA methylation measurement using targeted bisulfite sequencing [15]. We targeted CpGs that had the most significantly age-correlated methylation across the entire *Xenopus tropicalis* genome based on whole-genome bisulfite sequencing data from a previous study of 9 *Xenopus tropicalis* frogs [16]. Our approach captures *Xenopus*-specific age-associated methylation signals and is not restricted to sites that are conserved between amphibians and mammals. We therefore believe that our clock is a valuable tool for future studies in *Xenopus* aging biology and that our approach can be easily extended to other amphibians.

## Results

### Data Collection and Filtering

We collected DNA methylation data from 192 *Xenopus tropicalis* frogs with known ages from the National Xenopus Resource at the Marine Biological Laboratory (Woods Hole, MA, USA). The tissue samples used for DNA extraction were hindlimb webbing skin punches in the 187 adult frogs (Figure 1). For the 5 tadpoles, the entire organism was used for DNA extraction. Each frog was raised from birth in an aquatic tank with other frogs of the same age and strain. We collected samples from 26 distinct strains present across 35 physical tanks. The youngest animals in the dataset are 3-month-old tadpoles, and the oldest are 10.9-year-old adults. Supplementary Table 1 contains the age and strain information for each frog.

**Fig. 1.**
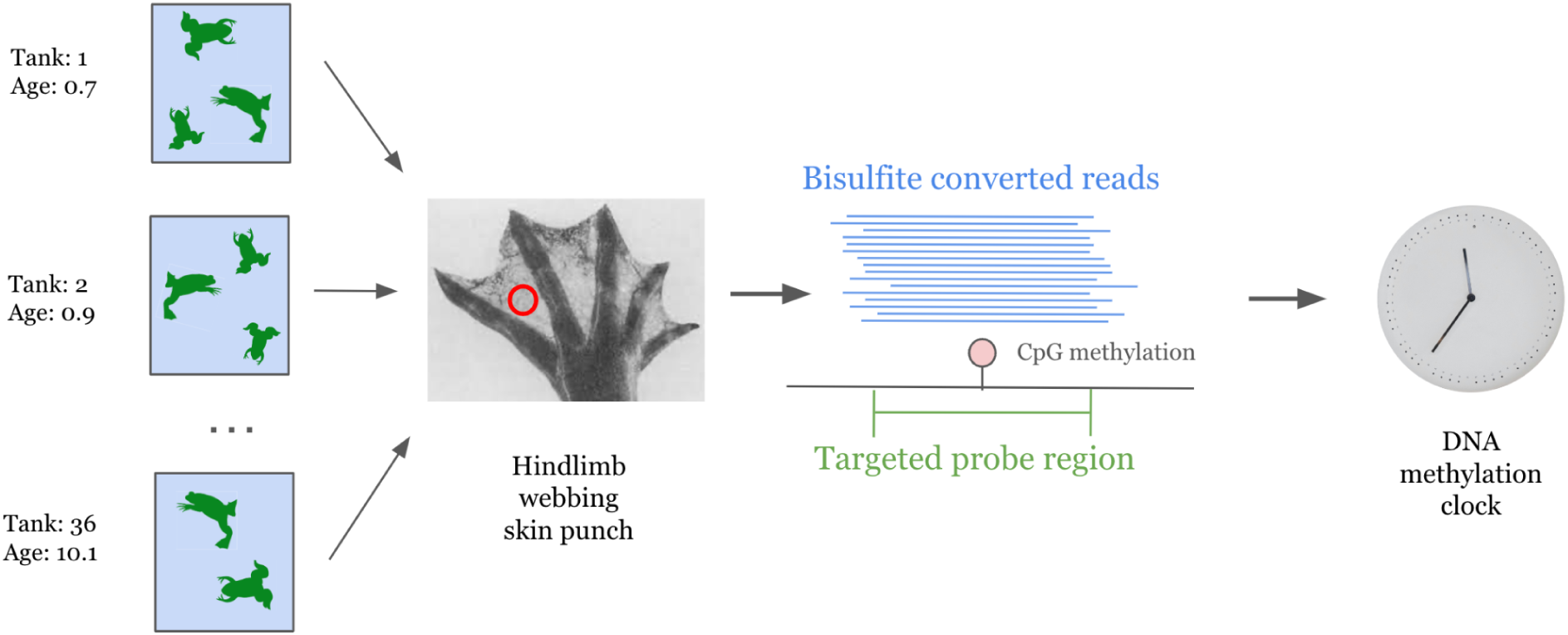
Overview of data collection workflow. *Xenopus tropicalis* frogs of different ages and strains were raised in tanks from birth, a skin punch from each frog’s hindlimb webbing was collected for DNA extraction, and targeted bisulfite sequencing was performed to measure DNA methylation levels at a collection of predetermined genomic regions. These targeted regions were designed to capture CpG sites that were previously shown to be highly age-associated in a genome-wide study of *Xenopus tropicalis*.

We used targeted bisulfite sequencing (TBS) to profile DNA methylation [15]. We designed DNA probes to cover regions of the genome that were previously shown to contain CpG sites with methylation highly correlated with age in our whole genome bisulfite sequencing (WGBS) study of 9 *Xenopus tropicalis* frogs of three different ages [16] (see Methods section). The 3,443 targeted probe regions are available in Supplementary Table 2. Bisulfite-converted reads underwent quality control, trimming, and alignment to the v10.0 reference genome [17] using BiSulfite Bolt [18] (see Methods section). In total, we measured 6,717 CpGs with sequencing coverage of at least 100x in all 192 frogs. Among the CpG sites shared between the whole genome and targeted bisulfite sequencing datasets, the age correlations calculated under the WGBS and TBS datasets have a significant positive correlation (Supplementary Figure 1). As expected, many sites in our TBS data display a strong positive or negative correlation with age (see two examples in Supplementary Figure 2A).

We noted that some CpG sites show a characteristic pattern with 3 separate levels of methylation across samples (Supplementary Figure 2B). This pattern of methylation is likely caused by germline C to T mutations at these genomic positions across some of our strains [19]. We identified these sites as those with methylation values that had a better fit in a Gaussian Mixture Model (GMM) with 3 components than a GMM with 1 (see Methods section). After filtering out these sites, 6,183 CpG sites remained for further analysis. Principal component analysis of our TBS methylation data for each frog shows that age is highly correlated with the first principal component, similar to the results found in WGBS data [16] (Spearman r = 0.739, Supplementary Figure 3A). The average methylation level across all CpGs for each frog in our data tends to decrease with age (Pearson r = -0.466, Supplementary Figure 3B).

To better characterize the genetic differences between frogs in our data, we called genetic variation from the targeted bisulfite sequencing data using BiSulfite Bolt. The matrix of 3,252 SNP differences is available in Supplementary Table 3. We calculated genetic variation as the number of SNP differences between frogs, divided by the total number of callable sites (1,003,455). Supplementary Figure 4A shows the average genetic variation between every pair of tanks. As expected, the intra-tank genetic variation tends to be smaller than the inter-tank genetic variation. Supplementary Figure 4B shows the genetic variation between individuals. While we found up to a maximum of 0.056% genetic variation between frogs in the dataset, our clocks are still able to accurately predict age across strains.

### Epigenetic Clock

We trained elastic net regularized linear regression models to predict chronological age based on DNA methylation levels. To evaluate this method, we performed cross-validation on the training dataset (Figure 2A-D), and trained a final model on all the training data (n=192) to predict the ages of 16 additional frogs in an external test set (n=16) (Supplementary Figure 5). In our dataset, all frogs within one tank are the same age and strain, so the leave-one-out cross-validation procedure (LOO-CV) (Figure 2A-B) includes frogs from the same tank in both the internal training and testing splits, which could lead to overfitting due to tank-specific features. To evaluate the model’s performance without a tank-related bias, we also performed leave-one-tank-out cross-validation (LOTO-CV) (Figure 2C-D). In this procedure, all frog samples from one tank are left out of the training data, then a clock is trained on all samples from the other tanks, and predictions are made on each frog in the left-out tank. To measure the performance of the clocks, we used the median absolute error between the predicted age and actual age (MedAE), and the Pearson correlation between the predicted age and actual age (r). As expected, the LOTO-CV predictions (MedAE = 0.835 years and r = 0.938) performed worse than LOO-CV predictions (MedAE = 0.456 years and r = 0.976). For comparison, the only other DNA methylation clock currently published in *Xenopus tropicalis* [12] achieved a LOO-CV MedAE of 0.96 years. We also report clocks that predict the natural logarithm of age (Fig 2B,2D), because previous work has suggested that DNA methylation levels tend to be non-linearly related to age [20,21]. The predictions of our log-transformed clocks show more accurate predictions for young samples and less accurate predictions for old samples.

**Fig. 2.**
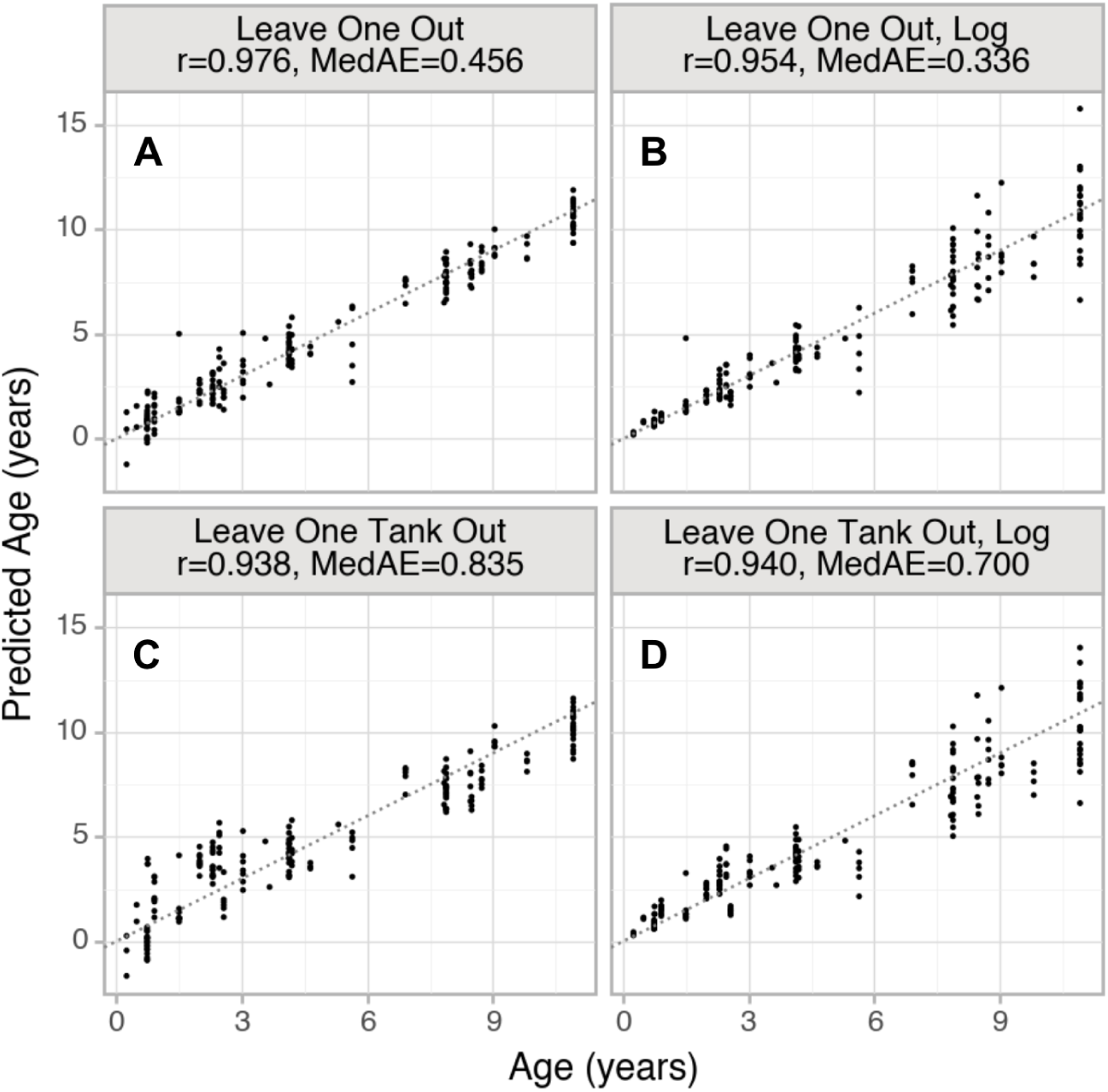
DNA methylation clock nested cross validation results on n=192 frogs. Models (B) and (D) were constructed to predict the natural logarithm of age. The dotted lines represent predicted age = actual age. MedAE is the median absolute error between predicted and actual age (in years).

For the elastic net model trained on the entire training dataset and evaluated on the 16 frog external test set, 378 CpG sites were selected as predictors (199 positively and 179 negatively associated with age). These CpG sites and the corresponding clock coefficients are provided in Supplementary Table 4. The external test set clock predictions (Supplementary Figure 5) display a very strong linear relationship between the predicted age and actual age (Pearson r = 0.991).

However, the slope of the predicted vs actual age plot is less than 1 (MedAE = 1.01 years). This reduction in the range of the predicted ages relative to the actual ages is a common feature of elastic net clocks when applied to validation data [22]. These 16 frogs were collected and processed separately from the 192 frog training data. However, it should be noted that some frogs in the test set came from the same tank and strain as the training data, so this evaluation shows generalization to different sample preparation and sequencing runs, but not necessarily generalization to entirely unseen frog strains and ages.

To further investigate the effects of including frogs of the same age and strain in both the training and prediction cross validation folds, we performed LOTO cross validation except with all the tank assignments randomized. This shuffled LOTO-CV procedure, when run 5 times with different tank membership assignments, achieved an average MedAE 0.470 years, and the log-transformed clock achieved MedAE of 0.398 years. The performance of the shuffled LOTO-CV is better than the original LOTO-CV, and similar to the LOO-CV results.

### Chromatin State Analysis

We sought to investigate whether highly age-correlated CpG sites are preferentially found in certain chromatin states. In the absence of an established chromatin state annotation for the *Xenopus tropicalis* v10.0 genome [17], we used ChromHMM [23] to construct one based on a previously published ChIP-seq dataset collected from *Xenopus tropicalis* embryos [24]. Our annotation is based on a stage 30 embryo from the published ChIP-seq data. The model generated 15 chromatin states which we assigned names based on previously reported chromatin states in other species. Supplementary Figure 6 shows the proportions of the top positively and negatively age-correlated CpGs in each of the annotated chromatin states.

Significantly age-correlated CpGs were selected based on adjusted p-values for the correlation between methylation and age (see Methods section). This generated a list of the top 326 positively age-correlated sites and 1907 negatively age-correlated sites. Most CpG sites in our data are found in the states corresponding to heterochromatin, quiescent, or repetitive regions of the genome. Figure 3A shows the emission probabilities output by ChromHMM. Figure 3B shows the log_2_ enrichment of the significantly age-correlated CpGs in each chromatin state, where enrichment is defined as the fraction of highly age-correlated CpG sites in a chromatin state divided by the fraction of all CpG sites in that chromatin state. We used Fisher’s exact test to compute a p-value for each enrichment. Although there are a small total number of CpGs in the “bivalent low” and “strongly transcribed” chromatin states in our dataset, these sites show the greatest enrichment for positively age-correlated CpGs. The heterochromatin1 state (moderate H3K9me3 signal) shows a moderate underrepresentation of highly age-correlated sites, and the heterochromatin2 state (strong H3K9me3 and H4K20me3, low H3K9me2) shows a moderate overrepresentation of highly age-correlated sites; these heterochromatin effects are significant under the Fisher’s exact test (Benjamini-Hochberg adjusted p < 10^-5^).

**Fig. 3.**
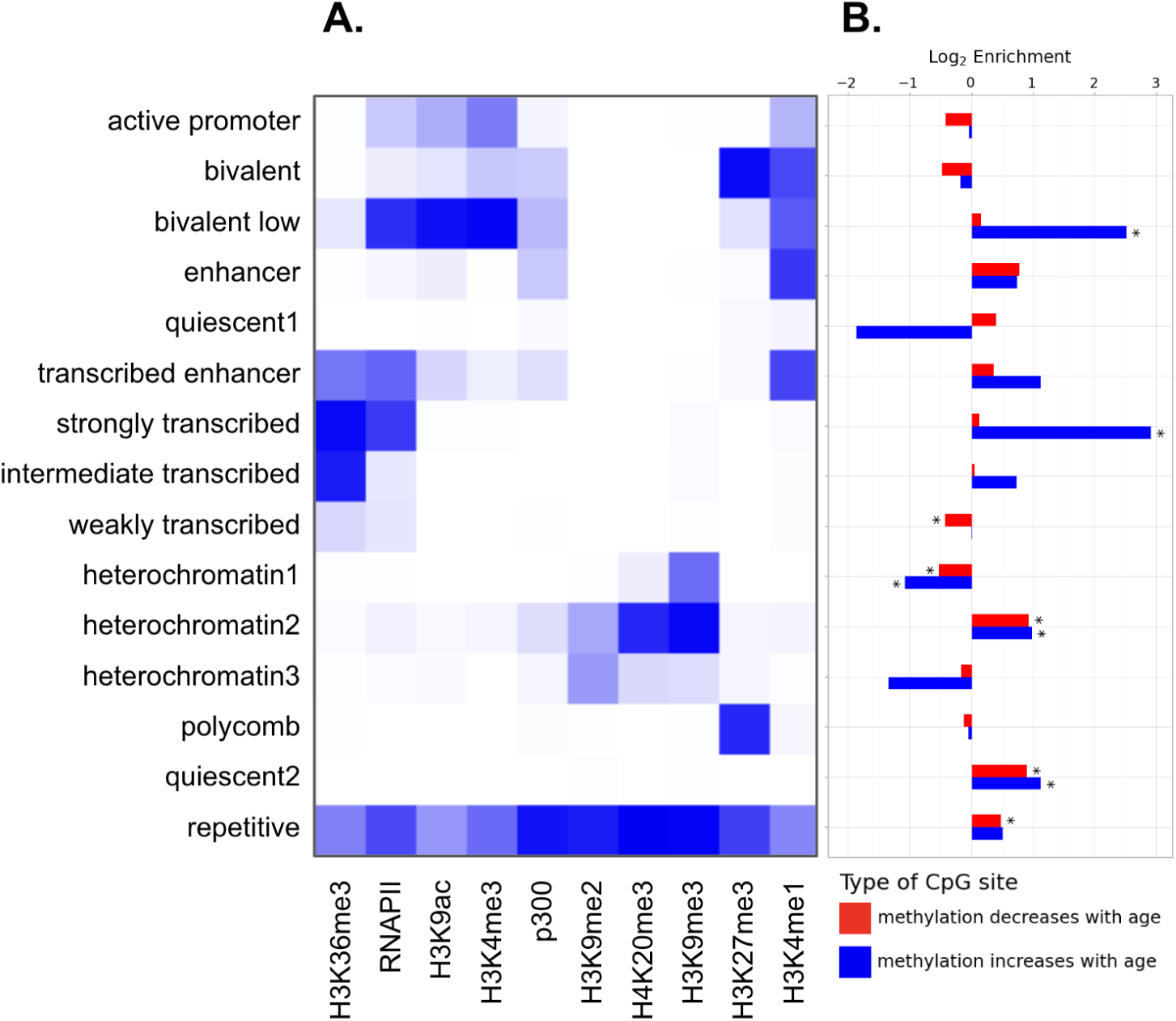
(A) ChromHMM automated chromatin state annotation emission parameters. (B) Chromatin states of selected CpG sites with high correlation between methylation and age. Log enrichment represents log_2_ of the fraction of + or - CpG sites in a chromatin state divided by the fraction of all CpG sites in that chromatin state. The reference background is all CpG sites from our TBS dataset with sequencing coverage of at least 10. An asterisk (*) indicates that the adjusted p-value < 0.05 for the Fisher’s exact test.

### Analysis of the expression levels of genes proximal to highly age-associated CpGs

We investigated the relationship between age-associated CpG sites and the expression of proximal genes. To accomplish this, we used the previously published *Xenopus* single-cell RNA-seq atlas [25]. The atlas includes gene expression data from 1-year-old frogs from the related species *Xenopus laevis*. For the purposes of this analysis, we mapped genes in *X. tropicalis* to *X. laevis*. We selected the significantly age-correlated CpGs in our data, and found the most proximal gene to each CpG site, generating a list of 37 positively- and 208 negatively age-correlated genes (see Methods section). Figure 4 shows the mean expression of these gene lists in each tissue, along with the mean expression of all genes in *Xenopus laevis* as a reference. The genes proximal to CpG sites with significant positive age association tend to be less expressed than the background of all genes, but due to the low sample size (37 positive genes), this effect is not significant under a Wilcoxon rank-sum test (Benjamini-Hochberg p-adj. > 0.05). The genes proximal to CpGs with significant negative age association are significantly more expressed than the background of all genes in brain, eye, heart, lung, muscle, ovary, pancreas, and stomach (Benjamini-Hochberg p-adj. < 0.05)

**Fig. 4.**
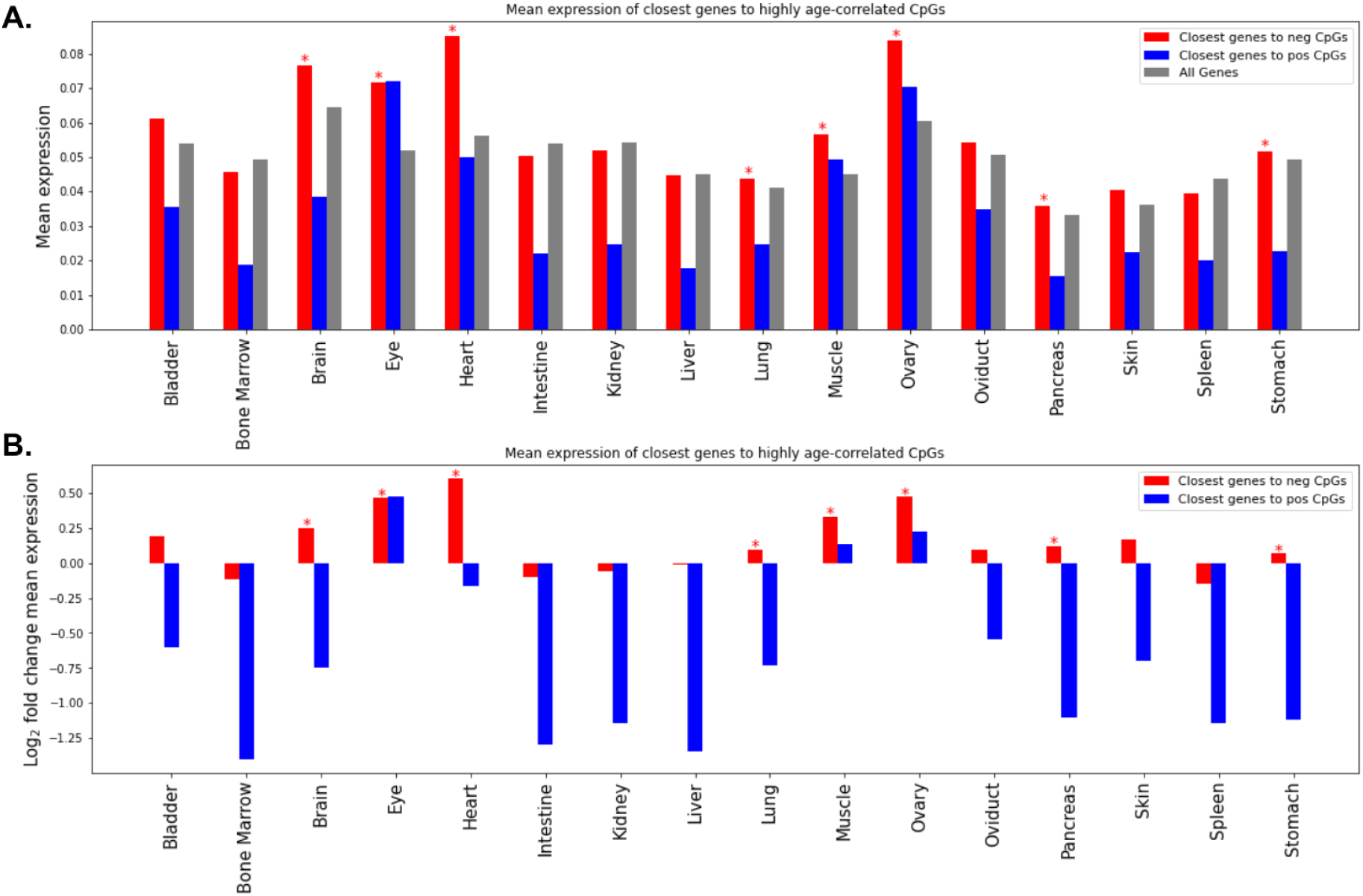
Single-cell RNA sequencing atlas expression levels of the closest gene to each significantly age-associated CpG site. Mean expression is calculated as the mean of log1pCP10k values over all cells in a tissue. * indicates Benjamini–Hochberg adjusted p-value is < 0.05 for Wilcoxon rank-sum test of mean expression in the gene list compared to mean expression of all genes.

## Methods

### Sample collection

192 Xenopus *tropicalis* frogs were housed at the National Xenopus Resource (RRID: SCR_013731) [26] in multi-rack recirculating aquatic systems with a constant temperature of 25°C and 12hr light /12hr dark cycle, and established diet as previously described [16,27,28]. Disposable biopsy punches were used to collect tissue (VWR 21,909–140) from the hindlimb webbing of each of the 187 adult frogs in the dataset [29]. For the 5 tadpoles, the entire organism was used for DNA extraction. Supplementary Table 1 includes all frog metadata, including strain. All animals have a Nigerian strain genetic background (NXR_1018). Some frogs in the dataset are transgenic; this information is given in the strain annotation. The sex of the frogs was not recorded. We performed DNA extraction according to the procedures described in [16], which was modified from [29].

### Targeted bisulfite sequencing library preparation

Library preparation was performed as previously described in [16]. 500 nanograms of purified DNA per sample was sonicated using a Covaris R230. Fragmented DNA was subject to end-repair, dA-tailing and ligation to pre-methylated unique-dual-indexed custom adapters (Integrated DNA Technologies). Pooled samples (16 samples per pool) were subject to hybridization with biotinylated oligonucleotide probes (synthesized by Integrated DNA Technologies) complementary to the previously identified age-associated regions (Supplementary Table 2) in the presence of *Xenopus* Hybloc DNA (Applied Genetics Laboratories - Melbourne, FL, US). Captured fragments were then treated with bisulfite (Zymo Lightning), then amplified and purified according to Morselli et al. [16]. Samples were subject to QC, normalized and combined in two pools of 96 samples, and each sequenced on NovaSeq6000 (SP lanes) as paired-end 150 bases.

### Targeted bisulfite sequencing data processing

The *X. tropicalis* v10.0 reference genome [17] was downloaded from Xenbase [30] (RRID:SCR_003280). We performed sequencing, alignment to the reference genome, quality control, and methylation calling using the same methods previously described in [16], except reads were trimmed using fastp [31] with TruSeq adapters trimmed and options --length_required 40 --trim_front1 5 --trim_tail1 5 --trim_front2 5 --trim_tail2 5 --trim_poly_g --qualified_quality_phred 20. Briefly, BiSulfite Bolt [18] was used to align bisulfite-converted reads to the reference genome and create a methylation matrix of all CpG sites with sequencing coverage of at least 100x in 100% of the samples. There are 6,717 of these CpGs. Mean coverage of each sample (available in Supplementary Table 1) was calculated with BSBolt AggregateMatrix (arguments: -count -min-coverage 100). We used scipy.stats.spearmanr [32] to compute the correlation between methylation and age at each site. For genotype analysis, we used the BiSulfite Bolt commands CallVariation and GenotypeMatrix to call variant genotypes from sites with a minimum of 10x coverage, alignment score 10, quality score 10, and present in at least 90% of frogs.

Some CpG sites in the data show an abnormal pattern of methylation, where some frogs have 0 methylation, some have 0.5 methylation, and some have 1.0 methylation, with very few values in between. We used the function sklearn.mixture.GaussianMixture [33] to fit Gaussian Mixture Models (GMM) to the methylation values at each CpG. A CpG site is filtered from the dataset if the Bayes Information Criterion (BIC) for the GMM with 1 component minus the BIC for the GMM with 3 components is greater than 90. The value of 90 was tuned manually to capture sites with clearly anomalous patterns. Additionally, for a site to be filtered there must be at least one vanishingly small methylation value (<0.006), since we expect the anomalous sites resulting from C to T germline mutation to have no methylation for frogs with the T allele. Examples of these sites are available in Supplementary Figure 1B. After filtering, our methylation matrix consisted of 6,183 CpG sites across 192 frogs.

Figures were generated with the Python packages plotnine v0.12.4 [34] and matplotlib v.3.8.2 [35]. Tabular data were manipulated using the Python packages pandas v2.1.4 [36] and anndata v0.8.0 [37].

### Epigenetic clock

We implemented the clocks using the function sklearn.linear_model.ElasticNetCV from the Scikit-learn package in Python [33]. The l1_ratio hyperparamter controls the relative strength of the L1 penalty compared to the L2 penalty, and it was fixed to 0.5 for the clocks. The alpha hyperparameter controls the overall strength of regularization. We used a nested cross validation approach to tune the alpha hyperparameter and obtain an estimate of the method’s performance. In this procedure, the n=192 frog training dataset is first split up into a training and validation set, based on the “outer cross validation” criterion of either LOO-CV or LOTO-CV. Within the training set of this cross validation, the ElasticNetCV function was used with an “inner cross validation” criterion to find the best alpha value within the training fold. For LOO-CV, the inner cross validation criterion was 5-fold cross validation. For LOTO-CV, the inner cross validation criterion was also LOTO-CV. Sklearn.model_selection.LeaveOneOut and sklearn.model_selection.LeaveOneGroupOut were used to implement these procedures.

Additionally, we used previously published (GEO Series GSE222107) [16] targeted bisulfite sequencing data from 16 different frogs as an external testing set for evaluating the performance of our DNA methylation clock. This external dataset was prepared by a different operator and sequenced on a different NovaSeq6000 lane. We trained a single clock on the subset of CpG sites shared between the main dataset and the test dataset (reducing the number of CpG sites from 6,183 to 5,938), then made predictions on the test dataset (Supplementary Figure 5). For this model, the function sklearn.linear_model.ElasticNetCV was used with internal LOTO-CV to tune the alpha hyperparameter.

### Selecting highly age-associated CpGs

From the 6,183 filtered CpG sites in our data, we selected the top sites most positively and negatively correlated with age for downstream analysis. We ranked the CpG sites based on their Spearman correlation between methylation value and age. For this site selection process, we included methylation measurements only from the 187 adult frogs and excluded tadpoles due to their substantially higher methylation values at some CpGs (see Supplementary Figure 3C). The p-value for this correlation was calculated with the Python function scipy.stats.spearmanr [32].

This p-value represents the null hypothesis that the Spearman correlation is 0. Highly age-associated sites were selected as those with a Benjamini–Hochberg adjusted p-value of less than 0.001. This approximately corresponds to a spearman r > 0.3 or r < -0.3. The result is 326 positively age correlated sites and 1907 negatively age correlated sites.

### Chromatin state analysis of age-associated CpGs

ChIP-seq data from Hontelez et al. were obtained from GEO Series GSE67974 [24]. The chromatin marks present in this dataset are H3K36me3, RNAPII, H3K9ac, H3K4me3, p300, H3K9me2, H4K20me3, H3K9me3, H3K9me3, H3K27me3, and H3K4me1. We used data from the stage 30 *Xenopus tropicalis* embryo. Cutadapt v2.10 (arguments: -u 5 -j 0) was used to trim the input FASTQ files [38]. Bowtie2 v2.4.2 [39] (arguments: -k 1 --no-unal) was used with bowtie2-build to align ChIP-seq reads to the v10.0 *X. tropicalis* reference genome. Samtools v1.11 view, sort, markdup, and index were used to process bam files and remove PCR duplicates [40]. ChromHMM v1.24 ConvertGeneTable, BinarizeBam, LearnModel, MakeSegmentation, and Reorder were used with default arguments to create the chromatin state annotation [23].

The number of chromatin states used in the genome segmentation is a user-controlled hyperparameter; we settled on 15 states to achieve a balance between having enough states to capture biological variation, and not so many to create duplicate states. We manually annotated the names of the 15 chromatin states based on the chromatin marks associated with each state. To statistically test CpG site enrichment for each chromatin state, we used Fisher’s exact test implemented with scipy.stats.fisher_exact, and adjusted the p values using the Benjamini-Hochberg method with stats.false_discovery_control [33].

### Expression levels of genes nearby to highly age-associated CpGs

First, we mapped each significantly age-associated CpG site in our dataset to the closest transcription start site (TSS) in the *X. tropicalis* v10.0 genome annotation [17] using pybedtools.closest v0.9.0 (argument: -D b) [41,42]. We excluded all CpGs whose closest gene TSS is greater than 5 kilobases away, since we wanted to capture proximal methylation effects on the nearby gene. Next, we mapped each *Xenopus tropicalis* gene to its homologous gene in the *Xenopus laevis* v9.1 genome [43] using a HMMER-based pipeline, as described in [44]. The 326 highly positively age-associated CpG sites in our *X. tropicalis* dataset mapped to 37 nearby genes with data available in the *X. laevis* atlas. The 1907 highly negatively age-associated CpG sites in our *X. tropicalis* dataset mapped to 208 nearby genes with data available in the *X. laevis* atlas. As a background reference, all available genes in the *X. laevis* expression dataset (25,362 genes) were used. First, the mean expression of each gene was computed across all cells of each tissue. Next, the mean expression over the genes in each of the three gene sets was calculated and displayed in Figure 4A. Figure 4B represents this same information by taking the log_2_ of the ratio between the expression of the positive or negative genes over the mean expression of all genes. The Python function scipy.stats.mannwhitneyu [32] was used within each tissue to test if the mean expression of the positive or negative gene sets are significantly different from the mean expression of all genes. scipy.stats.false_discovery_control [32] was used for the Benjamini-Hochberg p-value correction.

## Discussion

We constructed the most accurate clock for *Xenopus tropicalis*, based on 192 frogs. We expect the performance of this clock on new frog samples will be similar to the leave-one-tank-out cross-validation (LOTO-CV) performance metrics of MedAE=0.7 years and r=0.94. While the LOO-CV performance is better than LOTO-CV (MedAE=0.465 years, r=0.976), we were able to achieve similar performance to LOO-CV using a LOTO-CV approach with random tank membership assignments. This suggests that the LOO-CV procedure is overfitting to the age and/or shared genetic information from within each tank.

We identified chromatin states enriched for age-associated CpG sites using a chromatin state annotation based on a previously published ChIP-seq dataset in *Xenopus tropicalis* embryos [24]. The chromatin state primarily marked with H3K9me3 (“heterochromatin1”) makes up over 50% of the CpG sites in our dataset. We observed a significant underrepresentation of highly age-associated sites in the heterochromatin1 state. This state is likely a type of constitutive heterochromatin, and this result indicates methylation levels in these regions tend to be more stable with age than in other chromatin states. The “heterochromatin2” state is marked primarily with H3K9me3 and H4K20me3, and is significantly enriched for both significantly positive and negative age-associated CpG sites. The difference in the age-associated methylation behavior of these two chromatin states suggests a significant role for H4K20me3 in epigenetic aging. The histone marks H4K20me3 and H3K9me3 are directly linked to DNA methylation through the maintenance DNA methyltransferase DNMT1. DNMT1’s BAH1 domain has been shown to recognize H4K20me3 in humans [45], and the UHRF1 protein binds to H3K9me3 through its tandem Tudor domain and guides DNMT1 to these regions [46]. Therefore, changes of these histone marks with age could lead to age-associated DNA methylation changes at these loci. Large scale changes and redistribution of H3K9me3 and H4K20me3 have been observed with aging [47–49]. Since a large fraction of age-associated CpG sites in our dataset are in heterochromatic regions, it’s possible that much of our clock’s predictive ability can be explained by a redistribution of heterochromatin in *Xenopus* frogs with age. Direct measurement of the spatiotemporal evolution of these chromatin marks over aging is a possible future research direction.

We observed that positively age-correlated CpGs are significantly enriched in the state with H3K4me3 and H3K27me3 (“bivalent low”). This enrichment has been reported in human [10,50] and mammalian studies [11]. This effect has also been observed in *Xenopus tropicalis* and *laevis* [12] using chromatin state annotations for the bivalent sites from human datasets. Loss of H3K4me3 at chromatin that was formerly bivalent is a possible mechanism explaining the increase in DNA methylation with age. As H3K4me3 at bivalent sites is lost with age, we expect DNA methylation to increase at these sites because the *de novo* DNA methyltransferase machinery (DNMT3a/b/L) can bind to H3K4me0 (and not H3k4me3) through the ADD domain and promote *de novo* DNA methylation [51,52]. However, we note that less than 1% of CpG sites in our targeted bisulfite sequencing dataset are in this state (see Supplementary Figure 6). The targeted bisulfite sequencing method with different probes could allow for direct targeting of these sites in further studies.

The two chromatin states with the highest H3K27me3 levels (“polycomb” and “bivalent”) do not have significant enrichment for age-associated DNA methylation sites. This result contrasts with mammalian studies which report positively age-associated methylation at polycomb group target sites [10,53–56]. While a typical annotation of this chromatin state is defined as regions of the genome highly bound by PRC2 in embryonic stem cells, our annotation is based on measuring H3K27me3 in DNA from a whole *Xenopus* embryo.

We also observe significant enrichment for the top positively age-associated CpGs in chromatin with H3K36me3. H3K36me3 is found in gene bodies of actively transcribed genes, and maintenance of this mark prevents cryptic transcription [57]. DNMT3a’s PWWP domain interacts with H3K36me3, promoting *de novo* DNA methylation at these regions [58–60].

Similar to our results, DNA methylation levels have been found to increase with age in H3K36me3 regions in mouse muscle stem cells [61].

Using a single cell RNA-seq dataset from a young adult *Xenopus laevis* [25], we observed low expression of genes nearby to CpGs with positively age associated methylation. This effect has been previously observed in humans and mice [10,50,62–64]. The low expression levels of genes proximal to positively age-correlated CpGs suggest that the increase in methylation at these sites over age may generally not lead to significant phenotypic changes because these genes are already repressed by epigenetic mechanisms other than DNA methylation. This is consistent with the fact that positively age-associated CpGs have been found near developmental genes in humans [9,11] and *Xenopus* [12].

In conclusion, in this study we have shown that some mechanisms of epigenetic aging may be conserved between mammals and *Xenopus tropicalis*, supporting the future use of frogs as a model organism for the study of biological aging. An advantage of using amphibians in aging research is that they are cold blooded. The surrounding water temperature could be altered in a controlled experiment to determine the effect of metabolic rate on the rate and patterns of DNA methylation over time. Sexual maturity is well understood in *Xenopus*; specifically, metamorphosis can be easily controlled by iodine in the environment or by exogenous thyroid hormone. The delayed onset of sexual maturity increases lifespan in mice [65]. Further study of the molecular mechanisms of this process could be undertaken in *Xenopus*. While we did not consider the effect of sex in this study, future studies could examine lifespan differences between sexes using our clocks. Male-to-female and female-to-male sex reversal can be easily induced by exposing developing larvae to estrogen and the aromatase inhibitor fadrozole respectively [66]. There are also environmental factors that would be challenging to study in other animals, for example, the effects of environmental plastic exposure, which can be examined in *Xenopus* [67]. Moreover, *Xenopus* is thought to have negligible senescence and remains fertile late in life. In fact, it has been suggested that there is a lack of reproductive aging in *Xenopus* [68], which combined with our clocks could reveal new reproductive biology. It is known that wood frogs (*Lithobates sylvaticus*) are naturally freeze-tolerant [69]. If clocks translate to these species, exploring the clocks in frozen tissues presents a new research direction. Since single cell nuclear transfer was originally accomplished in *Rana* and *Xenopus* frogs [70], *Xenopus* provides a convenient toolbox for studying the germ line reset. In conclusion, *Xenopus* frogs provide promising research directions to further study the biology of aging.

## Acknowledgements

Supported by NIH’s R24 award OD031956 (L.P.). We sincerely thank Kseniya Petrova and William Ratzan (Harvard Medical School) respectively for technical assistance with *X. tropicalis* and *X. laevis* genome annotations, and preparation of *Xenopus* Hybloc DNA. We thank Silvia Nigro (University of Parma) for pre-processing bisulfite sequencing data. This project used computational and storage services associated with the Hoffman2 Cluster, operated by the UCLA Office of Advanced Research Computing’s Research Technology Group, and the High Performance Computing facility at the University of Parma (Parma, Italy). Frog samples were obtained from the National Xenopus Resource (RRID:SCR_01373).

## Data & Code Availability

Bisulfite sequencing FASTQ files and the processed methylation matrix for 192 frogs will be available at the Gene Expression Omnibus (GEO).

All code for the analysis will be available at github.com/ronanbennett/xenopus-aging.

## Supplementary information

Supplementary Table 1: *Xenopus* samples description

Supplementary Table 2: Targeted bisulfite sequencing probes

Supplementary Table 3: Genetic variation

Supplementary Table 4: Epigenetic clock weights

---

**Supplementary Figure 1:**
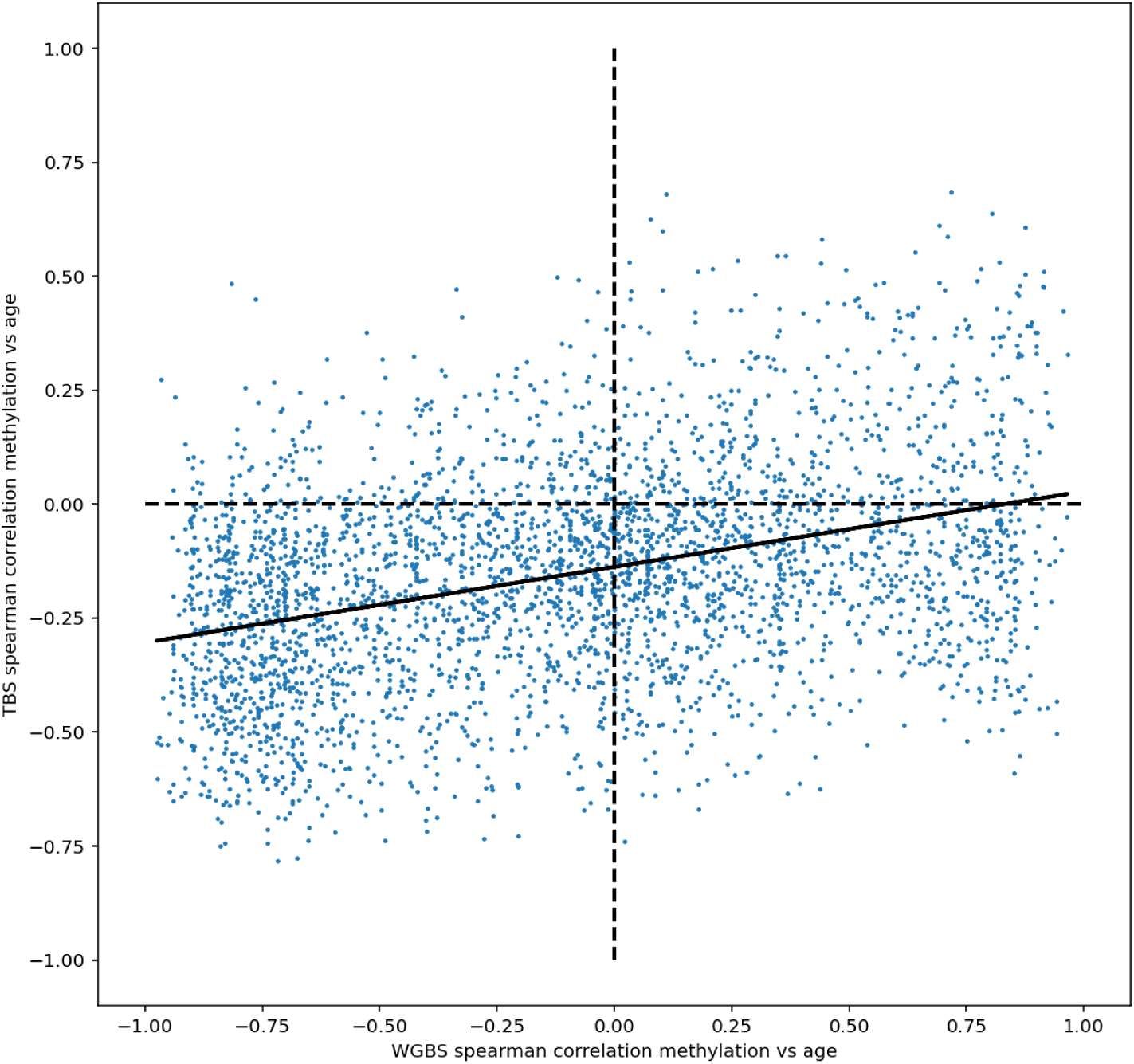
Comparison of n=3,506 CpG sites shared between the previously collected whole genome bisulfite sequencing (WGBS) dataset and this study’s targeted bisulfite sequencing (TBS) dataset. The Spearman correlation between methylation and age for the corresponding dataset is shown on each axis. The Spearman correlation between the Spearman correlations (WGBS vs TBS) in the plot is 0.307 (p<10^-100^) The solid black line shows the least squares regression best-fit line.

**Supplementary Figure 2:**
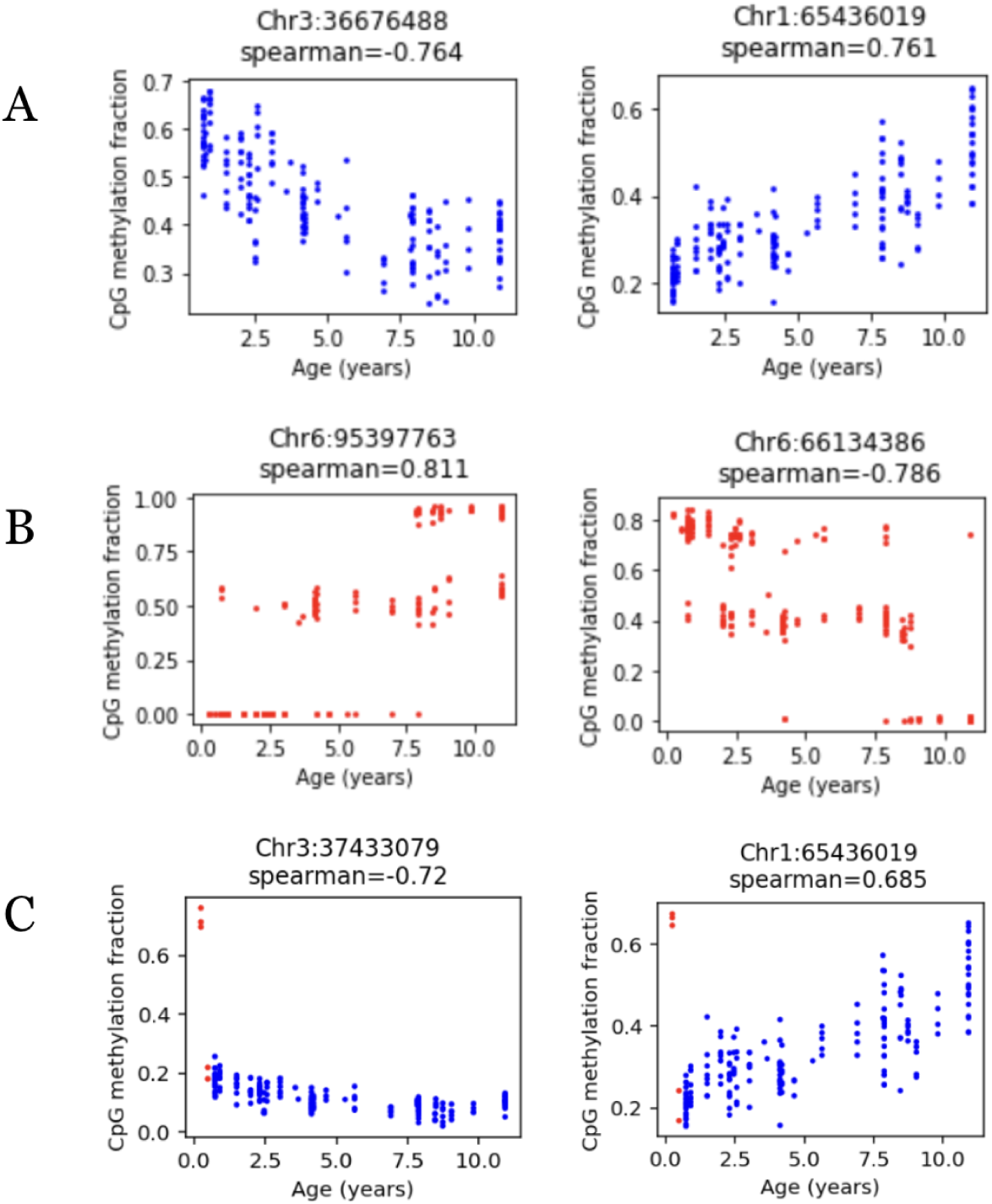
(A) Examples of CpG sites with highly age-correlated methylation. (B) Examples of CpG sites with a pattern of 3 distinct methylation levels. These are filtered out of downstream analyses. (C) Examples of CpG sites with abnormally high methylation levels in tadpoles (red). The tadpole samples are removed for some downstream analyses.

**Supplementary Figure 3:**
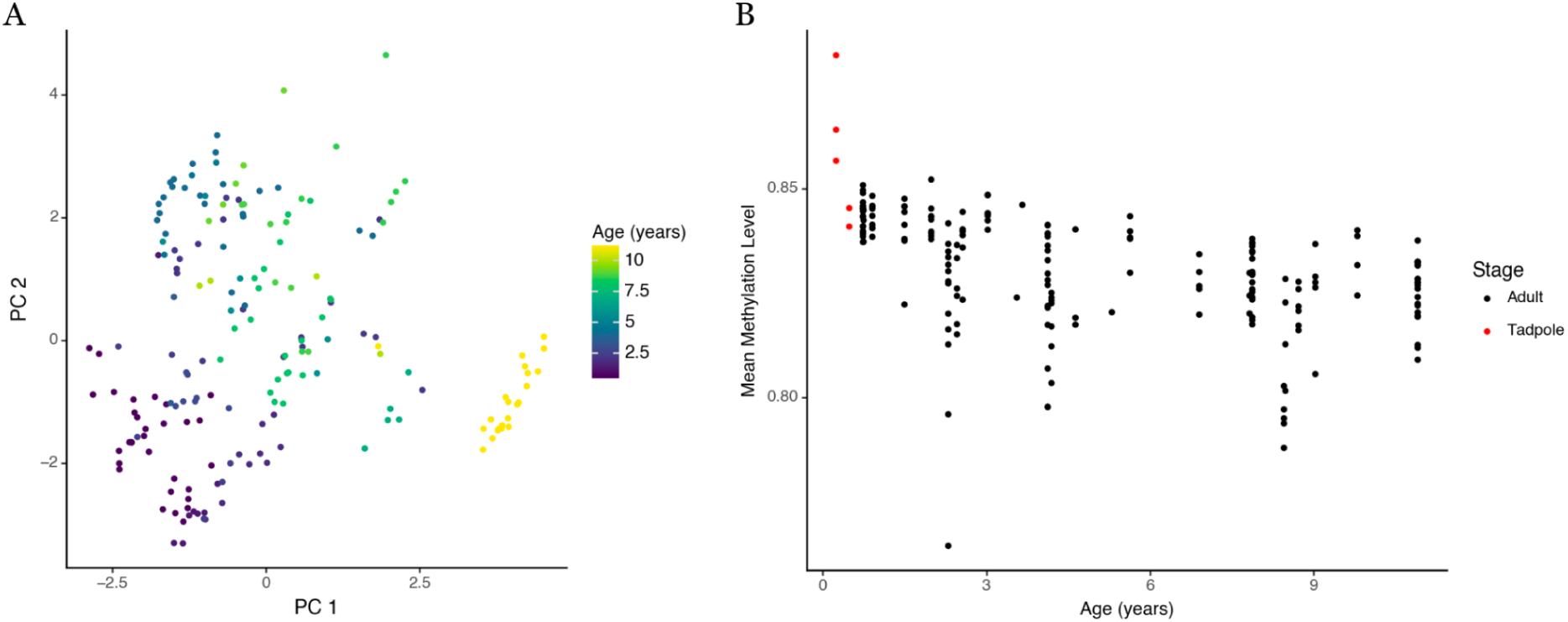
(A) Principal component analysis (PCA) of the DNA methylation values for each frog. Each CpG site’s methylation value was centered to the mean across all frogs before PCA. (B) Mean methylation level across all targeted bisulfite sequencing CpG sites vs age. CpG sites with a minimum coverage of 100 were used for both plots.

**Supplementary Figure 4:**
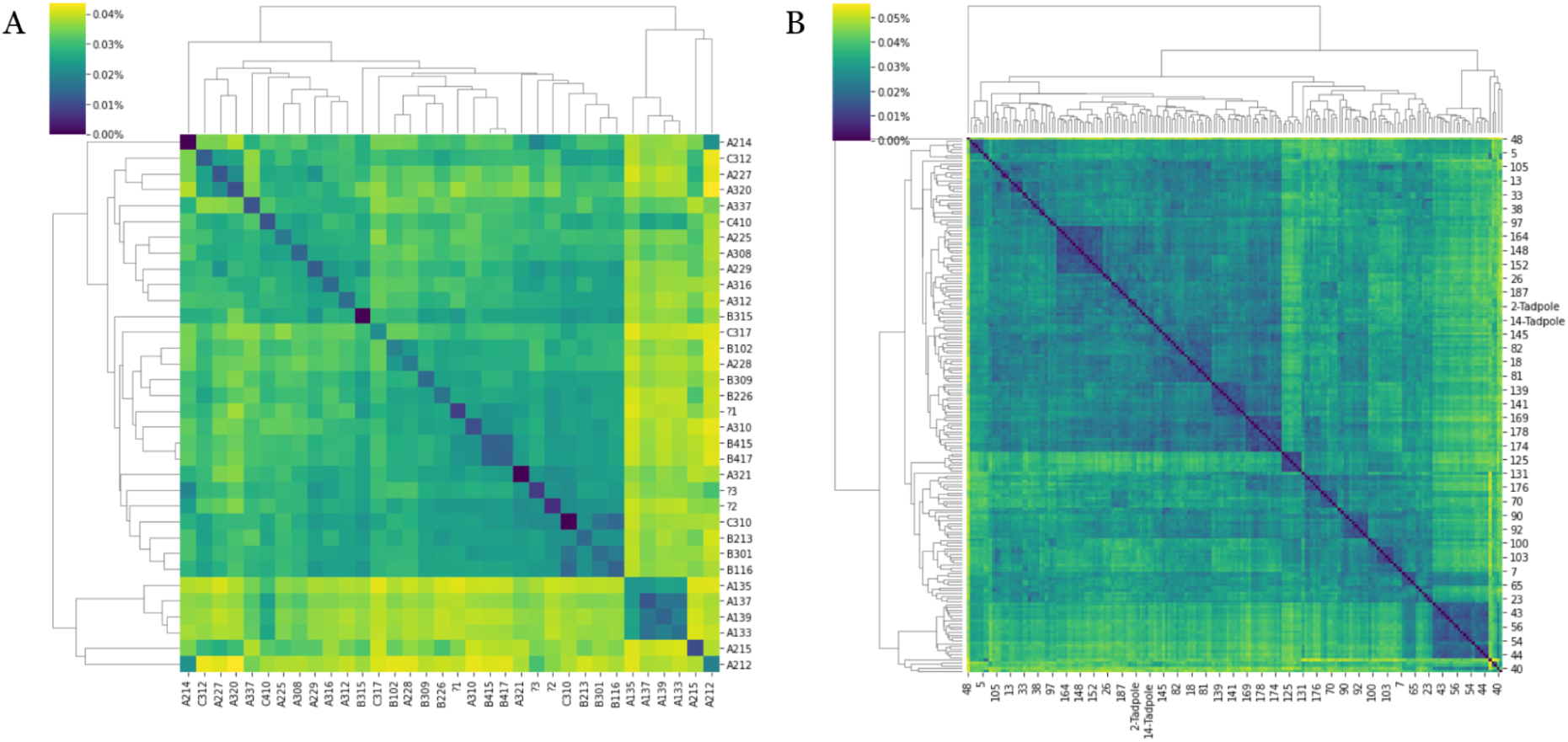
(A) Genetic variation between every pair of tanks (average number of SNP differences / number of callable base pairs). (B) Genetic variation between every pair of frogs (number of SNP differences / number of callable base pairs).

**Supplementary Figure 5:**
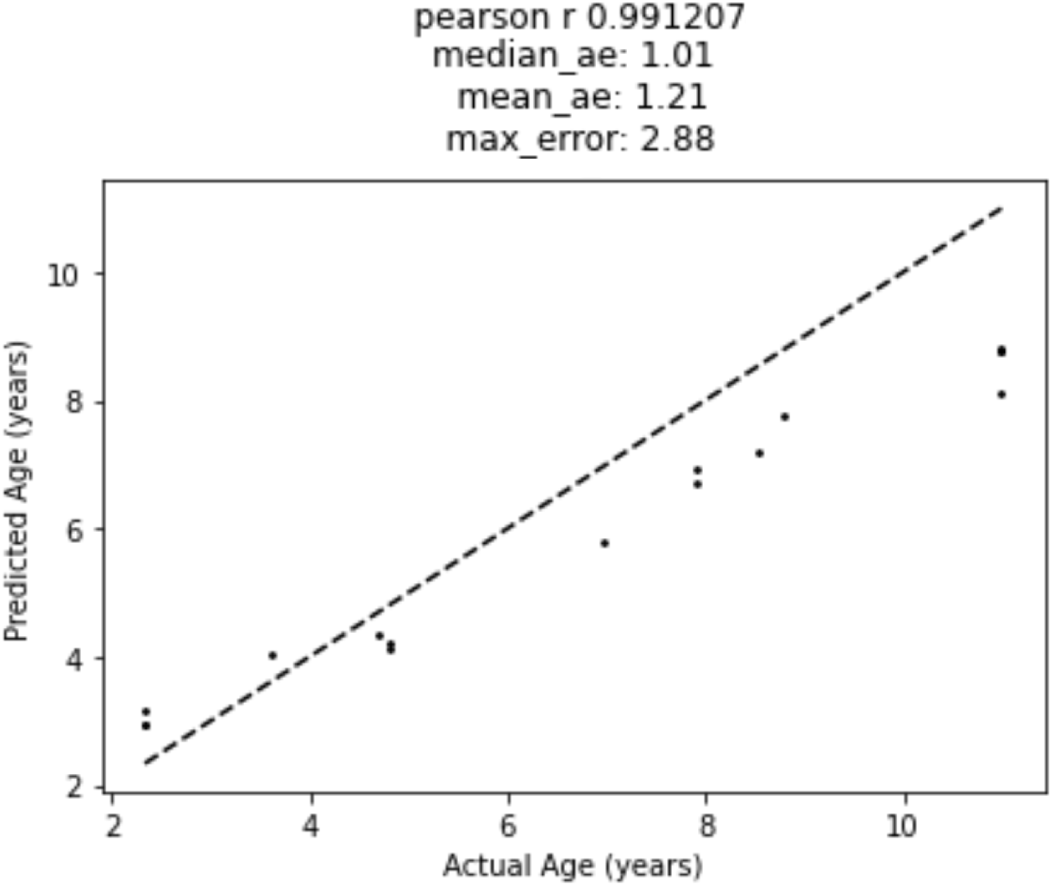
Age predictions on a test set of 16 frogs based on a DNA methylation clock trained on the 192 frog training set. The dotted line represents predicted age = actual age. The median absolute error, mean absolute error, and max error metrics are in units of years.

**Supplementary Figure 6:**
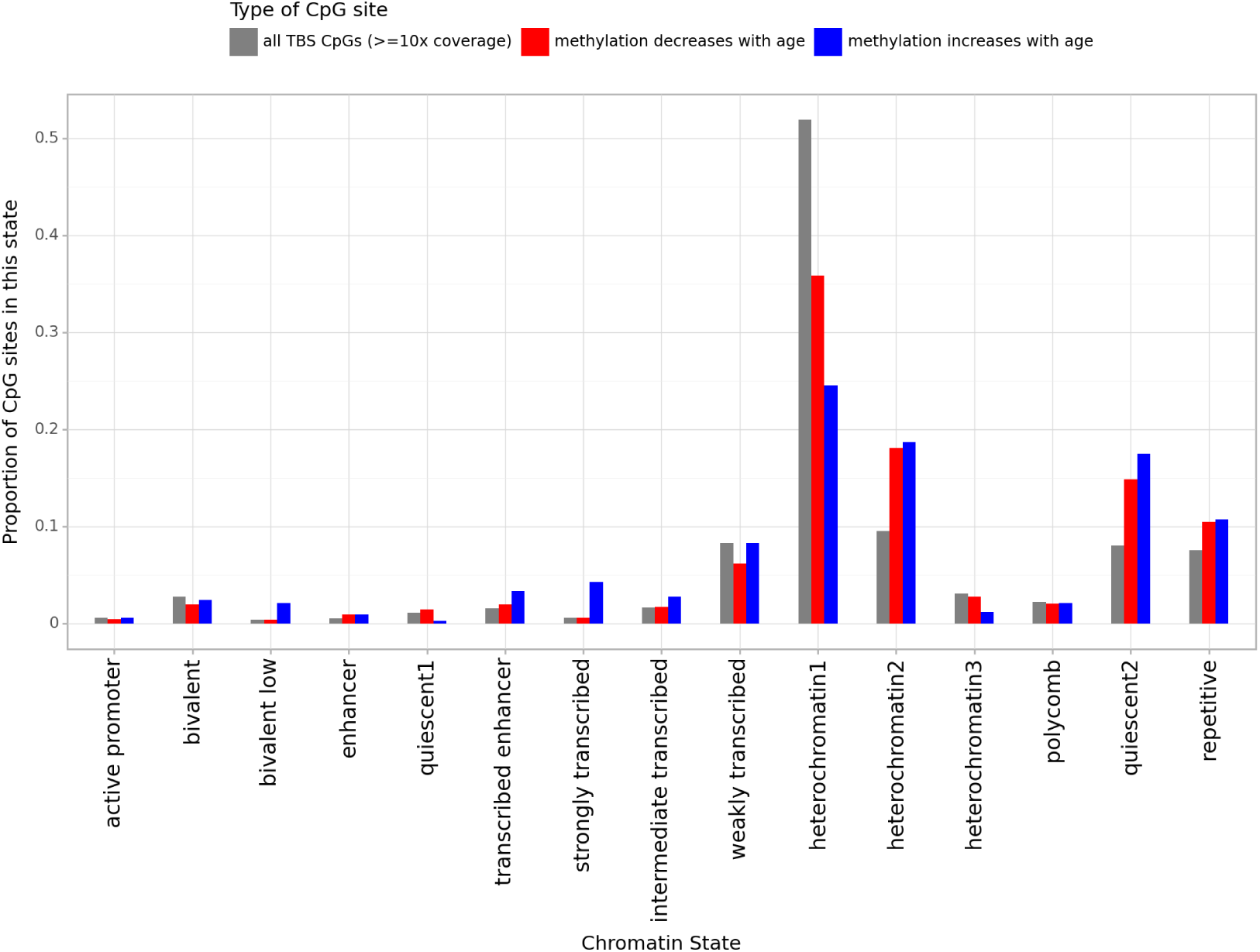
Proportion of CpG sites in each chromatin state for 3 groups of CpGs. There are 25,362 CpGs in the TBS dataset with >= 10x coverage (gray), 1,907 highly negatively age-associated CpGs (red), and 326 highly positively age-associated CpGs (blue).

